# Reduction in Ia afferent input via ischaemia alters motor unit discharge characteristics and estimates of persistent inward currents

**DOI:** 10.64898/2026.05.01.722246

**Authors:** Nikki Bonett, Tamara Valenčič, Christopher D Connelly, Haydn Thomason, Gregory EP Pearcey, Mathew Piasecki, Jakob Škarabot

## Abstract

Persistent inward currents (PICs) govern motoneuron output and are influenced by diffuse neuromodulation and local inhibition. When large diameter afferent feedback is lost, as in some neurological conditions, PICs might additionally amplify and prolong synaptic inputs. Here, we examined whether reducing Ia afferent transmission via ischaemic nerve block alters PIC contribution to tibialis anterior (TA) motor unit (MU) discharge.

Across two experiments 12 adults (5 female) performed triangular-shaped isometric dorsiflexion to 30% (Experiments 1 and 2) and 50% (Experiment 2) maximum voluntary force (MVF) at baseline, after a 20-minute rest (control), and during occlusion after inducing an ischaemic nerve block, confirmed by abolition of the soleus H-reflex. TA myoelectrical activity measured during contractions was decomposed into MU spike trains, and from smoothed MU discharges, discharge rate hysteresis (ΔF) and ascending non-linearity (brace height) were quantified.

Results from Experiment 1 involving contractions matched to absolute force levels revealed increased peak discharge rate, ΔF, and brace height post-occlusion. However, ΔF normalised to maximal theoretical hysteresis did not change across time points. In Experiment 2, where MVF was reassessed at each timepoint and contractions were matched to relative force, peak discharge rate, normalised ΔF and brace height increased post-occlusion compared to pre-, across both contraction intensities. ΔF only increased post-occlusion at 50% MVF, with no changes at 30% MVF.

These results show that ischaemic block of large-diameter axons, likely reducing reciprocal inhibition, increases PIC contribution to discharge rate modulation, highlighting the role of Ia afferent input in shaping motoneuron output in humans.

## INTRODUCTION

The force generated during voluntary contractions depends on both the number of motor units (MUs) recruited, and their discharge rate modulation (Heckman & Enoka, 2012). The neural activation that a MU transforms into mechanical and contractile activity is impacted by numerous inputs, which converge on the α-motoneuron (Burke et al., 1992). These inputs can be broadly categorised into ionotropic inputs (e.g. corticospinal, reticulospinal, propriospinal, and afferent inputs), and metabotropic input from the brainstem (also known as neuromodulation; (Heckman et al., 2009; Kuo et al., 2003). Brainstem neuromodulation, mediated by monoaminergic input (e.g. serotonin [5-HT] and noradrenaline [NE]), facilitates persistent inward currents (PICs) on motoneuron dendrites (Heckman et al., 2005; Lee & Heckman, 2000) via voltage-sensitive calcium (Carlin et al., 2000) and sodium (Lee & Heckman, 2001) currents. PICs provide gain control of the motor system (Lee & Heckman, 1996) and introduce non-linear discharge behaviour of motoneurons (Heckman et al., 2008; Johnson et al., 2017), characterised by amplification and prolongation of motoneuron discharge in response to linear increases and decreases in excitatory synaptic input, allowing for MU derecruitment at a lower threshold relative to recruitment (i.e., hysteresis; Heckman et al., 2005).

Although 5-HT and NE facilitate PICs by controlling the motoneuron’s intrinsic excitability and sensitivity to ionotropic input (Heckman & Enoka, 2012), their effects are diffuse, affecting multiple motoneuron pools simultaneously (Heckman et al., 2005; Lee & Heckman, 2000). In the case of excitation, while muscle co-activation or sustained muscle activation is sometimes necessary, a control mechanism which can better specify motor pool activation when this becomes excessive is essential (Heckman & Enoka, 2012; Pearcey et al., 2022). Inhibition (e.g. via peripheral afferent input) is the most likely control mechanism, sculpting local motor pool excitability from the diffuse monoaminergic neuromodulatory environment (Heckman & Enoka, 2012). Indeed, in the decerebrate cat, a stretch of the antagonist muscle inducing a small increase in reciprocal inhibition has been shown to result in ∼50% reduction in dendritic PICs (Hyngstrom et al., 2007), suggesting that PICs are highly sensitive to inhibition (Hultborn et al., 2003; Kuo et al., 2003). In human studies, Vandenberk & Kalmar (2014) demonstrated a negative relationship between estimated reciprocal inhibition (via common peroneal nerve stimulation) and PIC contribution to self-sustained discharge rate (via discharge rate hysteresis of motor unit pairs; ΔF). Furthermore, increasing reciprocal inhibition via common peroneal nerve stimulation (Mesquita et al., 2022), tendon vibration (Orssatto et al., 2022; Pearcey et al., 2022), or exaggerated co-contraction (Gomes et al., 2024) all reduce MU discharge rate hysteresis and/or changes in MU discharge patterns consistent with reduced estimates of PICs. Nevertheless, contradictory evidence has also been demonstrated in humans where estimates of PICs have typically been shown to be greater at shorter muscle lengths (Beauchamp et al., 2025; Goreau et al., 2025; Valenčič et al., 2026) despite an expectation for reciprocal inhibition to be greater compared to longer lengths.

To date, however, no human studies have demonstrated a link between *reduced* reciprocal inhibition and its effect on estimates of PICs, which likely reflects the experimental difficulty in downregulating reciprocal inhibition, independent of changes in voluntary drive or co-activation. Understanding how reduced Ia afferent input modulates PICs has clinical relevance, given that loss of afferent input has been implicated in motoneuron hyperexcitability and, therefore, spasticity following stroke and spinal cord injury (Elbasiouny et al., 2010; Mazzaro et al., 2007; Takeoka & Arber, 2019). Prolonged ischaemic occlusion induces a nerve block, which presents a viable solution to the methodological challenge of downregulating reciprocal inhibition. Ischaemia preferentially blocks large-diameter afferents (including Ia) before motor axons, due to their greater metabolic vulnerability to hypoxia (Hofmeijer et al., 2013). The selectivity of ischaemic block via occlusion has been confirmed by abolition of the H-reflex concomitant with preservation of the maximal M-wave, and has thus been used in studies with the aim of isolating the contribution of Ia afferents to voluntary motor output (Lorentzen et al., 2018; Rasul et al., 2022). By blocking Ia afferents from the antagonist muscles, ischaemic occlusion provides a means of assessing how reduced disynaptic reciprocal inhibitory drive onto agonist motoneurons (Crone et al., 1987) influences PIC contribution to MU discharge rate in humans.

Therefore, the aim of this study was to investigate how reducing Ia afferent input via ischaemia impacts tibialis anterior (TA) MU discharge patterns during isometric dorsiflexion. We performed two experiments to compare discharge rate characteristics during isometric dorsiflexion 1) at the same *absolute* contraction intensity before and after occlusion, and 2) at the same *relative* contraction intensity to determine whether the potential changes in discharge rate characteristics were dependent on contraction intensity. It was hypothesized that decreased Ia afferent input would downregulate reciprocal inhibition and increase the estimates of PIC-related contributions to discharge, leading to increased discharge rate hysteresis and greater ascending discharge rate non-linearity.

## MATERIALS AND METHODS

### Participants

This study consisted of two experiments. Of the fifteen participants recruited for Experiment 1, three were unable to complete the post-occlusion measurements due to the inability to match the target force levels. Therefore, both experiments consisted of 12 young, recreationally active adults (Experiment 1: 5 females, 24 ± 4 years, 1.69 ± 0.10 m, 67.0 ± 13.0 kg; Experiment 2: 5 females, 25 ± 4 years, 1.72 ± 0.13 m, 69.5 ± 16.2 kg). Participants had no known neurological conditions or musculoskeletal injuries affecting their lower limbs, and were not taking any medication that could affect the nervous system. Before taking part, participants provided written informed consent, and their eligibility was assessed using a health screening questionnaire. The study procedures were approved by the Loughborough University Ethical Advisory Committee and the University of Nottingham Faculty of Medicine and Health Science Research Ethics Committee, and were conducted in accordance with the Declaration of Helsinki, except for registration in database.

### Experimental Design and Protocol

#### Experiment 1

Participants attended two laboratory sessions within 14 days. The first visit served as a familiarisation session where participants became accustomed to the experimental procedures by practicing triangular-shaped isometric dorsiflexion to 30% of maximum voluntary force (MVF). They were also habituated to percutaneous tibial nerve stimulation. Twenty-four and 12 hours before the second experimental session, participants were asked to avoid strenuous lower-body exercise and caffeine, respectively.

During the experimental session, participants were seated in a custom-built seat equipped with an ankle ergometer (for details, see below). Following preparation of skin and placement of EMG electrodes, participants were instructed to warm up their dorsiflexors by performing unilateral submaximal isometric dorsiflexion (3 × 50%, 3 × 75%, and 1 × 90% of perceived MVF). After the warm-up, participants performed two maximal effort ankle dorsiflexion with verbal encouragement (>30 seconds rest in between). If the two trials differed by more than 5%, a third maximal effort dorsiflexion contraction was performed. The highest instantaneous value was taken as MVF, and used for normalisation of force recordings.

We used a multiple baseline, repeated measures design. In the initial stage of the experiment (PRE1), participants performed a 20-second triangular-shaped isometric ramp contraction up to 30% MVF, composed of 10-second linear ascending and descending phases. To facilitate accuracy in ramp contraction performance, real-time feedback was provided on a computer screen ∼1.5 metres in front of participants. After the performance of triangular contractions, participants were provided with a 20-minute rest period (approximately equivalent in duration to the average occlusion period) during which they were taken out of the rig and seated on a chair. Once back in the rig, they performed the triangular isometric ramp contraction to 30% MVF once again, as a second baseline measure (PRE2).

Electrical stimuli were then delivered to the posterior tibial nerve to evoke Hoffman reflexes (H-reflexes) in the soleus. First, an H-reflex/M-wave (H/M) recruitment curve up to maximal M-wave (M_max_) stimulation was evoked. Following that, stimulation intensity was reduced to induce an H-reflex on the ascending limb of the H/M curve with a preceding M-wave amplitude of 5-10% M_max_. M-wave amplitude was kept consistent to ensure the same proportion of the motor pool was activated each time a reflex was evoked (Zehr, 2002). A 10 cm sphygmomanometer cuff was then fitted above the knee and was, using the Hokanson Rapid Cuff Inflation System (Hokanson, Bellevue, WA, United States), immediately inflated to 200 mmHg to induce an ischaemic nerve block (Nielsen et al., 1992). H-reflex amplitudes were monitored every 2 minutes until they dropped to below 10% of the original value (Grey et al., 2001; Lorentzen et al., 2018). In practice this resulted in the near-complete abolishment of the H-reflex (see Figure 1A for example response). Once the H-reflex was abolished, a triangular isometric ramp contraction to 30% MVF was repeated before a final M_max_ was evoked, the inflation system was turned off, and the cuff was deflated.

**Figure 1.**
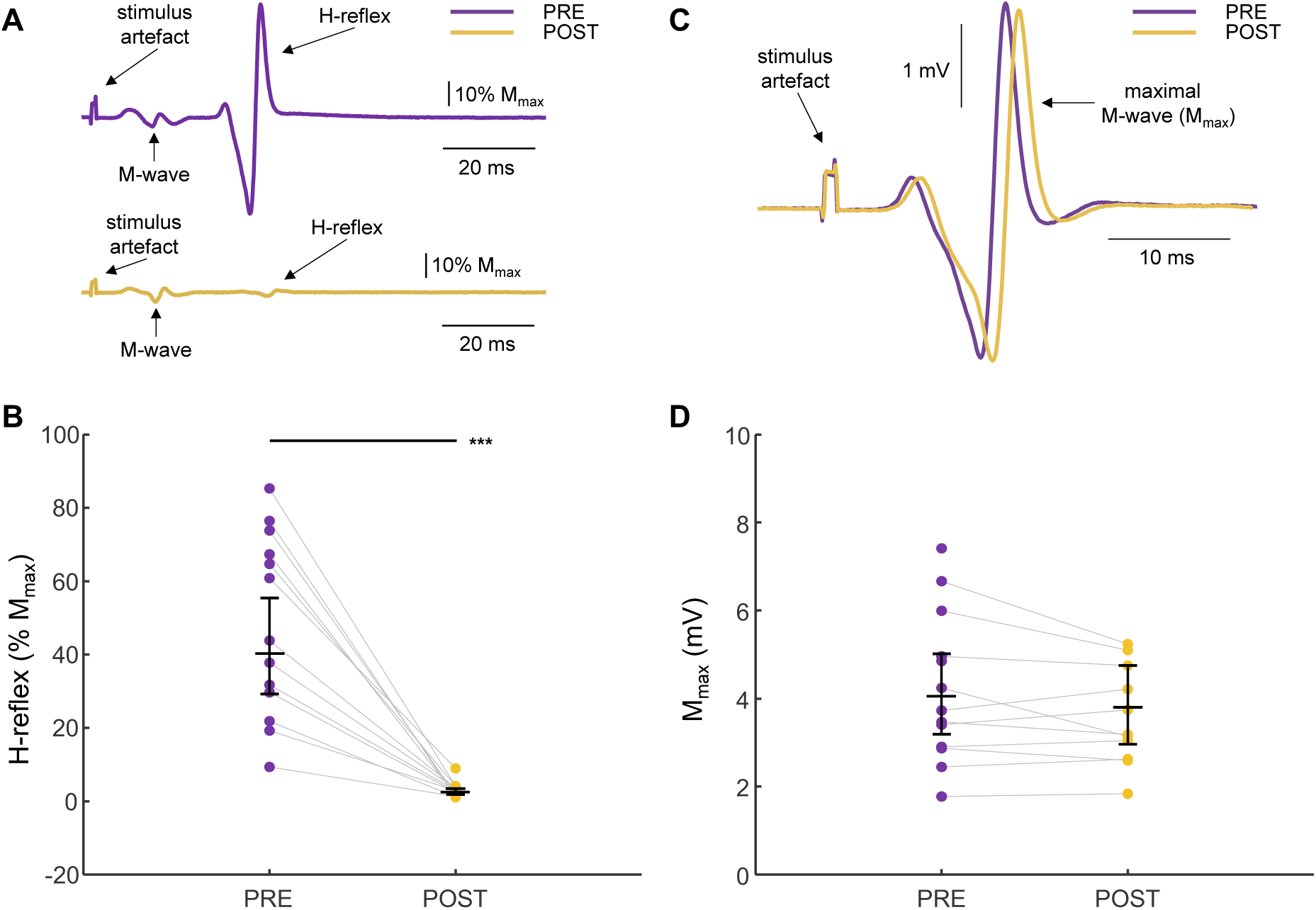
Soleus H-reflex and maximal M-wave before and after occlusion. **A.** Example soleus H-reflex in a representative participant at PRE (before occlusion using a sphygmomanometer cuff inflated to 200 mmHg above the knee) and at POST (once H-reflex was almost completely abolished). **B.** Filled circles represent mean H-reflex amplitude as a percentage of maximal M-wave (% M_max_) before (PRE) and after (POST) occlusion for each participant. The mean value at each time point was calculated by taking the average H-reflex amplitude in response to 3 stimulations delivered at least 5 seconds apart during that condition. **C.** Example M_max_ traces from a representative participant before occlusion (PRE) and at the point when H-reflex is near complete abolishment at the very end of the experiment (POST). **D.** Filled circles represent M_max_ amplitude in millivolts (mV) before (PRE) and after (POST) occlusion for each participant. Note the absence of one connecting line due to a missing data point for one individual at POST. For each condition in B. and D., the middle horizontal black bar represents the estimated marginal mean obtained from linear mixed modelling with error bars denoting the 95% confidence intervals. *** p < 0.001.

#### Experiment 2

In Experiment 1, triangular contractions were performed to the same absolute force level before and after occlusion. However, it has previously been shown that occlusion might affect maximal force-producing capabilities of the agonist muscle after occlusion (Glencross & Oldfield, 1975). In order to account for this potential reduction in MVF, Experiment 2 was conducted in a similar way to Experiment 1, but with the addition of a warm-up before PRE2 and MVF contractions performed at each time point (PRE1, PRE2, and POST). Ramp contractions were then performed to the same relative contraction intensity. In order to further explore the potential contraction intensity-dependent effects of occlusion, triangular contractions were performed to both 30% and 50% MVF, based on MVF at each time point (PRE1, PRE2, and POST).

### Experimental Procedures

#### Force Recordings

Participants sat on a custom-made chair with their foot in an ankle ergometer equipped with a force transducer (CCT Transducers s.a.s., Torino, Italy). Participants were positioned with hips flexed at 110°, knee fully extended (180°) and ankle at 10° of plantar flexion. One foot was secured to the ergometer at the metatarsophalangeal joints using two straps: one made of 35 mm-wide reinforced canvas webbing, and one made of Velcro. The analogue signals from the force transducer were amplified (×200; Forza-B; OT Bioelettronica, Torino, Italy), and simultaneously sampled in Spike2 (version 10; Cambridge Electronics Design Ltd., Cambridge, UK) and OTBiolab+ (OT Bioelettronica, Torino, Italy) software using an analogue-to-digital converter (ADC; Micro 1401-4, Cambridge Electronics Design Ltd., Cambridge, UK) and a 16-bit multichannel amplifier (Quattrocento; OT Bioelettronica, Torino, Italy), respectively.

#### High-Density Surface Electromyography

To record HD-sEMG signals from the TA, a semi-disposable 64-electrode grid (13 rows × 5 columns, 1 mm electrode diameter, 8 mm inter-electrode distance; GR08MM1305, OT Bioelettronica, Torino, Italy) was used. The grid was aligned with a disposable bi-adhesive foam layer (SpecsMedica, Battipaglia, Italy), the cavities of which were filled with conductive paste (SpecsMedica, Battipaglia, Italy). After skin preparation involving shaving and abrasion, the grid was adhered to the skin over the TA muscle belly. A reference electrode (Kendall Medi-Trace, Cardinal Health, Dublin, OH, USA) was adhered to the medial malleolus and a dampened strap ground electrode was wrapped around the ankle of the contralateral limb. The monopolar HD-sEMG signals were bandpass filtered (10-500 Hz), sampled at 2048 Hz using the Quattrocento 16-bit multichannel amplifier, and acquired using OTBiolab+ software.

#### Limb Occlusion

Following PRE2, a sphygmomanometer cuff (SC10, 11 × 85 cm cuff size, 10 × 41 cm bladder size; Hokanson, Bellevue, WA, United States) was instantly inflated to 200 mmHg above the knee using a rapid cuff inflation system (Hokanson, Bellevue, WA, United States). The limb occlusion period varied in length for each participant, which lasted the combined duration of time it took for the abolishment of H-reflex and to perform contractions at the POST timepoint. The cuff remained inflated during the final ramp contraction at POST in Experiment 1, and during the recording of an MVF contraction and the ramp contractions to 30% and 50% MVF at POST in Experiment 2. The cuff was deflated immediately after the final M_max_ was evoked.

#### Tibial Nerve Stimulation

Percutaneous electrical nerve stimulation (1 ms pulse duration; DS7R, Digitimer Ltd., Welwyn Garden City, UK) was performed to elicit H-reflexes and M-waves in the soleus. A bipolar stimulating electrode (Digitimer, Hertfordshire, UK) was placed over the tibial nerve in the popliteal fossa and its position was adjusted until a strong H-reflex was evident in the bipolar soleus EMG signal recorded in Spike2 software. Following correct participant positioning, H/M recruitment curves were obtained and M_max_ was elicited in the soleus, to enable response normalisation. M_max_ was obtained by gradually increasing stimulus intensity until the amplitude of the EMG response plateaued, after which the stimulus intensity was further increased by 25% to ensure supramaximal stimulation. Stimulus intensity was then decreased, and H-reflex was elicited on the ascending limb of the H/M curve, with the M-wave amplitude preceding H-reflex maintained at 5-10% M_max_. Once the cuff was inflated, H-reflex was monitored every 2 minutes until it was abolished (<10% H-reflex amplitude from PRE) through the delivery of 3 stimulations (≥5-second interstimulus interval to avoid post-activation depression; Burke, 2016). M-wave amplitudes were monitored and kept constant at 5-10% M_max_ amplitude to ensure constant stimulation delivery to the nerve (Pearcey & Zehr, 2020). After performing triangular-shaped isometric dorsiflexion to 30% MVF in the POST condition in Experiment 1, and triangular isometric dorsiflexion to 30% and 50% MVF in Experiment 2, a final M_max_ was elicited.

### Data Processing and Analysis

#### H-Reflex and M-wave Analysis

Offline analysis of responses to tibial nerve stimulation in soleus was performed using Spike2 software. Peak-to-peak amplitude of M_max_ was recorded at the point before occlusion commenced (PRE) and just before the sphygmomanometer cuff was deflated (POST). The peak-to-peak amplitude of three soleus H-reflexes recorded every 2 minutes during the occlusion process were also calculated.

#### High-Density Surface Electromyography

The HD-sEMG signals from the triangular-shaped isometric dorsiflexion contractions collected throughout the two experiments were analysed in MATLAB using customised scripts (R2023b; The Math-Works, Matick, MA). Signals were digitally bandpass filtered (10-500 Hz) using a zero-lag fourth-order Butterworth filter and decomposed into individual MU spike trains using a Convolution Kernel Compensation algorithm (Holobar & Zazula, 2007). The MU spike trains were manually edited using established procedures (Murks et al., 2025), and those exhibiting low pulse-to-noise ratio (<28 dB; Holobar et al., 2014) were excluded.

#### Motor Unit Discharge Characteristics

To understand changes in MU discharge patterns with occlusion, instantaneous MU discharge rates were smoothed using support vector regression with a Gaussian kernel (kernel scale of 1.6, box constraint of 370) and the adaptive epsilon parameter (interquartile range/11; Beauchamp et al., 2022). Recruitment threshold was calculated as the force at the instant of the first spike in the MU spike train, and peak discharge rate was taken to be the maximal MU discharge taken from the smoothed estimate of MU discharge rate.

As previously indicated, estimates of the effects of PICs on MU discharge can be ascertained from the MU discharge patterns. PIC-related prolongation of MU discharge leads to MU derecruitment at a lower synaptic input than at recruitment. We quantified prolongation by estimating onset-offset hysteresis of a lower-threshold (reporter) MU at the onset and offset of a higher-threshold (test) MU (ΔF; Gorassini et al., 1998, 2002). The pair of MUs was considered suitable based on three criteria: 1) to ensure full activation of the PIC in the reporter unit, the pair of MUs must have had a recruitment time difference greater than 1 second (Bennett et al., 2001; Hassan et al., 2020; Powers et al., 2008); 2) to ensure the MU pairs received common synaptic input, they must have had rate-rate correlations of r^2^ > 0.7 (Gorassini et al., 2004; Stephenson & Maluf, 2011); and 3) to minimise the influence of MU saturation on the calculation of ΔF, the reporter unit must have had a discharge rate greater than 0.5 pps while the test unit was active (Stephenson & Maluf, 2011). Because multiple reporter units may be suitable for comparison with each test unit, a mean of all ΔF values for a given test unit was calculated (Hassan et al., 2021). Additionally, to account for variations in descending discharge rate modulation, ΔF were normalised to the maximal theoretical hysteresis for each test-reporter unit pair; i.e., the difference in reporter unit discharge rate at test unit recruitment and reporter unit’s discharge rate at derecruitment (Škarabot et al., 2025).

The contribution of PICs to the ascending discharge rate modulation was quantified as the ascending discharge rate non-linearity (known as brace height; Beauchamp et al., 2023). Specifically, the maximum orthogonal vector relative to a theoretical line between the points of initial and peak discharge rate was calculated. This value was additionally normalised to the height of the right triangle with a hypotenuse between the points of initial and peak discharge rate (percentage of the right triangle; % rTRI).

### Statistical Analysis

All statistical analyses were performed in R software (version 4.5.1, R Foundation for Statistical Computing, Vienna, Austria) using RStudio (version 2025.05.1+513, RStudio Inc., Boston, MA, USA). For each outcome variable, modelling was performed independently, and histograms and quantile-quantile plots of model residuals were used to assess distribution.

Linear mixed models were constructed for MU-related metrics with participant as random effect and time point (PRE1, PRE2 or POST), contraction intensity (30% MVF and 50% MVF; for Experiment 2 only) and their interaction as fixed effects. Additionally, as decomposition is biased towards higher-threshold MUs (Farina et al., 2010), recruitment threshold was included as a covariate. Data points corresponding to residuals falling outside ±3 z-scores were deemed to be outliers and removed (Experiment 1: peak discharge rate = 18 outliers, ΔF = 7 outliers, normalised ΔF = 4 outliers, brace height = 3 outliers; Experiment 2: peak discharge rate = 10 outliers, ΔF = 8 outliers, normalised ΔF = 3 outliers, brace height = 2 outliers). If data distribution was skewed, data were logarithmically transformed or square rooted; the type of transformation was selected based on the fit of the model. The residuals of predictions were found to be non-linear for the following variables: peak discharge rate in Experiment 1 (log-transformed), normalised ΔF in Experiment 1 (sqrt-transformed), normalised ΔF in Experiment 2 (sqrt-transformed) and brace height in Experiment 2 (sqrt-transformed).

The significance of the models was assessed by comparing the fit of the model with and without predictors using the *lmerTest* package (Kuznetsova et al., 2017). Main effects and interactions were assessed via Analysis of Variance using Wald chi-square statistics. Using the *emmeans* package, pairwise post hoc tests of estimated marginal means were performed on the significant main effects and interactions and were adjusted using the Tukey method (Lenth, 2025). Where data had been transformed, results from post hoc tests were back-transformed to present data on the original scale, and effect sizes were calculated as standardised mean differences (Cohen’s d). The latter were computed by dividing the estimated marginal mean difference by the residual standard deviation of the linear mixed model. Linear mixed models were also constructed for M_max_ amplitude (data sqrt-transformed in both Experiments 1 and 2) and H-reflex amplitude (data log-transformed in both Experiments 1 and 2), with participant as random effect and time point (PRE or POST) as fixed effect to assess whether M_max_ differed across timepoints and to verify that H-reflexes were abolished by means of occlusion. Significance was set at α = 0.05, and results are presented as estimated marginal means [95% confidence intervals].

## RESULTS

### Experiment 1

#### H-reflex and M-wave Analysis

During the period of occlusion (mean ± standard deviation of occlusion duration: 17 ± 7 min), peak-to-peak amplitude of the soleus H-reflex was measured as a percentage of M_max_ amplitude, on the ascending limb of the recruitment curve, whilst preceding M-wave amplitude was maintained at 5-10% M_max_. When H-reflexes were abolished, ischaemic nerve block was deemed effective (for example reflexes, see Figure 1A). Indeed, H-reflex amplitude changed from PRE to POST (Χ^2^_(1)_ = 798.88, p < 0.0001), being lower at POST (2.50 [1.82, 3.44]% M_max_) compared to PRE (40.30 [29.28, 55.43]% M_max_, p < 0.0001, d = 6.40; Figure 1B), indicating Ia afferent input was sufficiently reduced. To ensure that peripheral nerve fibre membrane excitability was not altered by occlusion, soleus M_max_ peak-to-peak amplitude was calculated before and after occlusion with an example response shown in Figure 1C. The soleus M_max_ was not significantly different between timepoints (Χ^2^_(1)_ = 2.55, p = 0.1101; 4.05 [3.19, 5.02] vs 3.80 [2.96, 4.75] mV, p = 0.1345, d = −0.65; Figure 1D), confirming peripheral nerve fibre membrane excitability did not differ as a result of occlusion.

#### Motor Unit Discharge Characteristics

To appreciate the qualitative differences in MU discharge patterns before and after occlusion, ensemble averages were constructed (Beauchamp et al., 2022) and are shown in Figure 2A. Peak discharge rate changed across the experimental time points (Χ^2^_(2)_ = 207.98, p < 0.0001), having been found to be greater in the POST condition (25.4 [22.4, 28.9] pps) compared to PRE1 (19.8 [17.4, 22.5] pps, p < 0.0001, d = 1.38; Figure 2B) and PRE2 (20.7 [18.2, 23.5] pps, p < 0.0001, d = 1.14). Peak discharge rate also increased in PRE2 relative to PRE1 (p = 0.0321, d = 0.24), however, the magnitude of this effect was small.

**Figure 2.**
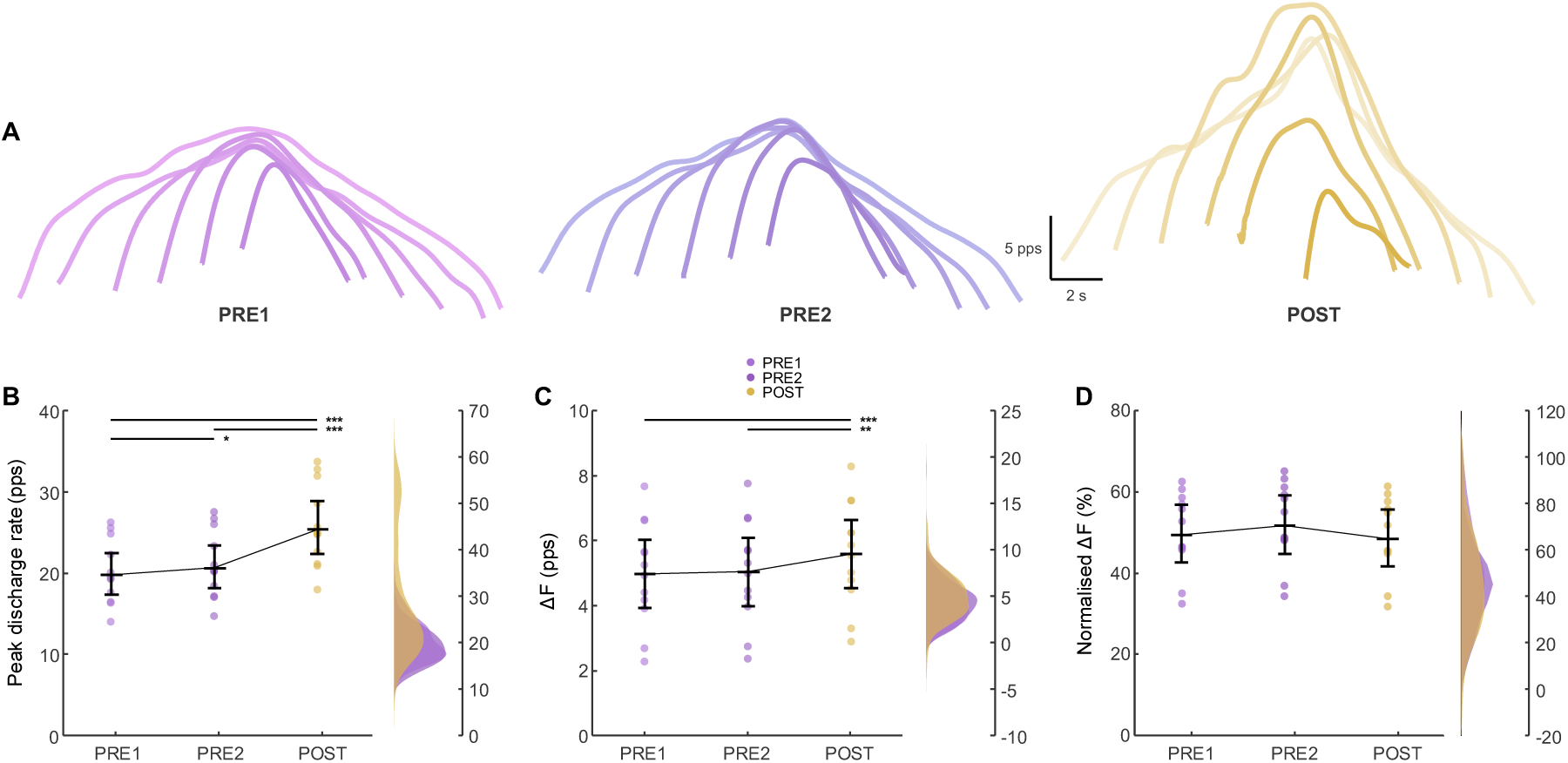
Motor unit discharge characteristics before and after occlusion. **A.** Ensemble smoothed motor unit discharge patterns at baseline (PRE1) after a 20-minute rest period (PRE2), and after occlusion (POST). **B.** Peak discharge rate in pulses per second (pps), **C.** discharge rate hysteresis (ΔF) in pulses per second (pps), **D.** ΔF normalised to the maximal theoretical hysteresis (%) at PRE1, PRE2 and POST. In B., C., and D. filled circles on the left-hand side of each figure represent individual participant averages over the three conditions. The middle horizontal black bars represent the estimated marginal means obtained from linear mixed effects modelling with error bars denoting the 95% confidence intervals. The kernel density distribution plots on the right-hand side of the figures show the distribution of points across values of peak discharge rate, ΔF, and normalised ΔF, respectively. *** p < 0.001, ** p < 0.01, * p < 0.05.

Onset-offset hysteresis also changed across the time points (Χ^2^ = 16.28, p = 0.0003), with ΔF found to be greater at POST (5.60 [4.55, 6.66] pps) compared to PRE1 (4.99 [3.93, 6.04] pps, p = 0.0007, d = 0.44) and PRE2 (5.06 [4.01, 6.11] pps, p = 0.0023, d = 0.39; Figure 2C). However, no differences in ΔF were observed between PRE1 and PRE2 (p = 0.8801, d = −0.05). Conversely, when ΔF was normalised to the maximal theoretical hysteresis, it did not appear to vary across time points (Χ^2^ = 3.03, p = 0.2201; Figure 2D).

To estimate the ascending MU discharge rate non-linearity, geometric analyses were performed where MU discharge rate is expressed as a function of dorsiflexion force; the ensemble averages for each timepoint are shown in Figure 3A. The ascending MU discharge non-linearity, measured via brace height, changed across the three conditions (Χ^2^_(2)_ = 18.27, p = 0.0001). While it was found to be greater at POST (42.4 [37.7, 47.0]%) compared to PRE1 (35.6 [31.2, 39.9]%, p = 0.0004, d = 0.60; Figure 3B), brace height at POST was not found to be significantly different to PRE2 (40.2 [35.9, 44.5]%, p = 0.4244, d = 0.19). Additionally, at PRE2, brace height was found to be greater than at PRE1 (p = 0.0042, d = 0.41).

**Figure 3.**
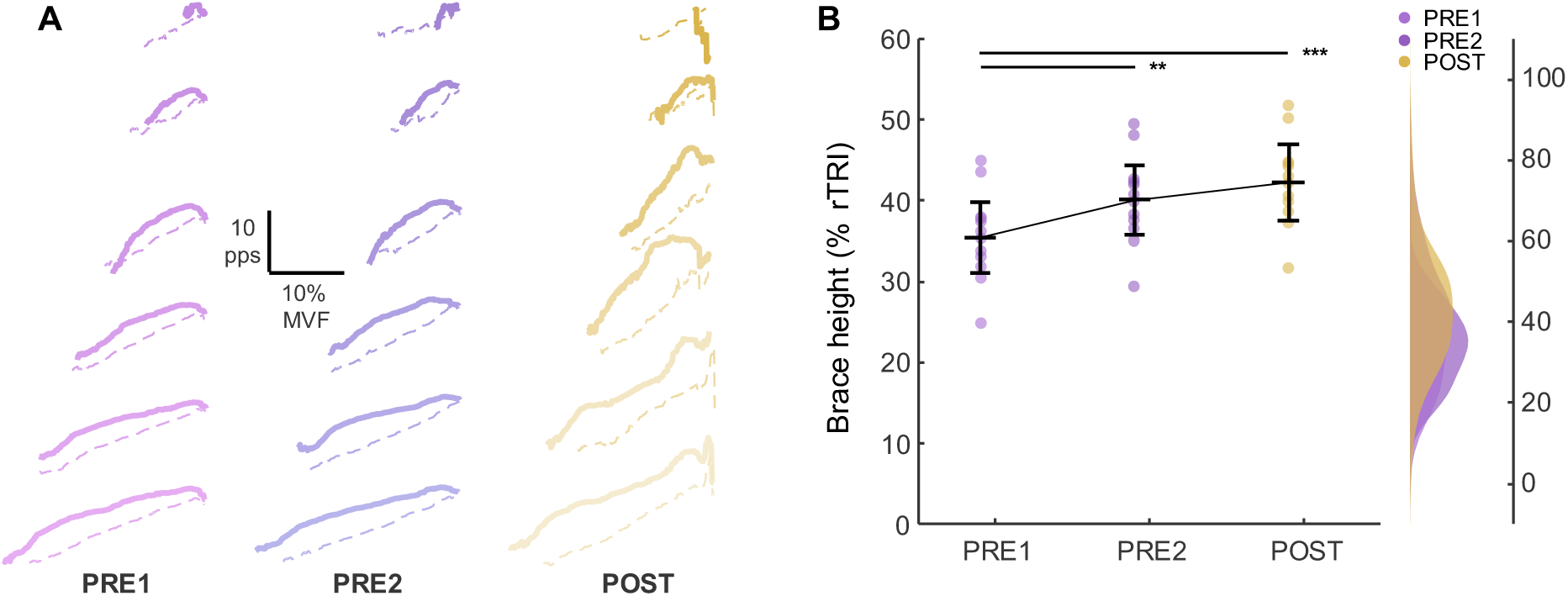
Geometric analyses of motor unit discharge rate before and after occlusion. **A.** Ensemble smoothed motor unit discharge patterns as a function of force (as % of maximal voluntary force, %MVF) during ascending (full lines) and descending (dashed lines) phase of triangular contractions at baseline (PRE1), after a 20-minute rest period (PRE2), and after occlusion (POST). **B.** Brace height as a percentage of a right triangle (% rTRI) at PRE1, PRE2, and POST. In B., filled circles on the left-hand side of the figure represent individual participant averages over the three conditions. The middle horizontal black bars represent the estimated marginal means obtained from linear mixed modelling with error bars denoting the 95% confidence intervals. The kernel density distribution plots on the right-hand side of the figures show the distribution of points across values of brace height. *** p < 0.001, ** p < 0.01.

### Experiment 2

#### Maximum Voluntary Force

Experiment 2 was conducted to account for the potential reduction in MVF post-occlusion, meaning participants could perform ramp contractions to 30% and 50% of their MVF at each time point. Indeed, we found a difference in MVF across time points (Χ^2^_(2)_ = 75.55, p < 0.0001). As expected, there was a decrease in MVF post-occlusion (172 [113, 244]N) compared to at PRE1 (283 [205, 374] N, p < 0.0001, d = 3.06) and PRE2 (284 [206, 375] N, p < 0.0001, d = 3.09), with no difference between PRE1 and PRE2 (p = 0.9980, d = - 0.02; Figure 4A). Despite POST contractions being performed to MVF at that time point, one participant was unable to perform contractions at 50% MVF post-occlusion, resulting in unequal sample sizes across contraction intensities, which was accommodated by the use of linear mixed-effects models.

**Figure 4.**
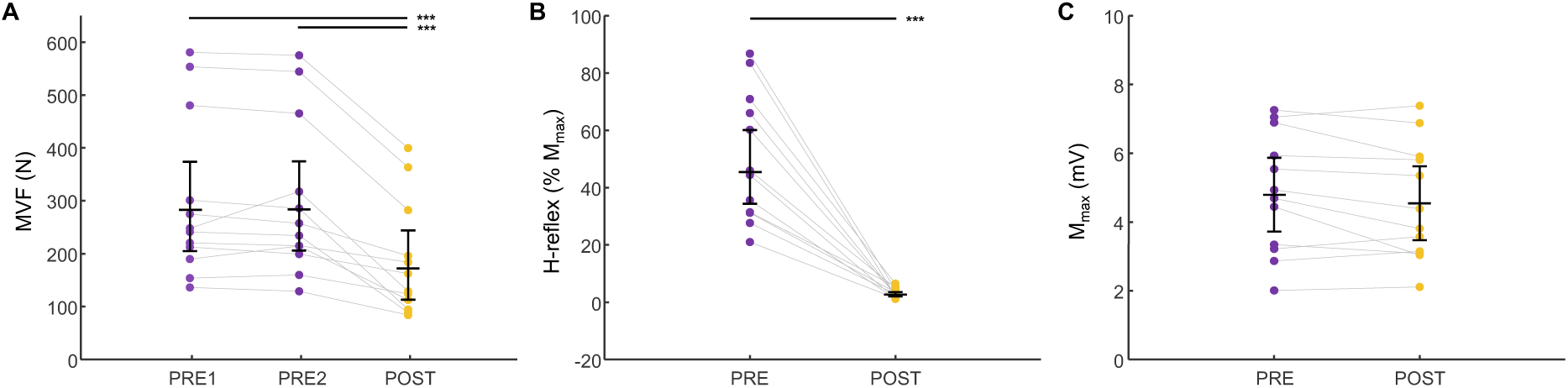
Maximal voluntary force, soleus H-reflex and maximal M-wave. **A.** Filled circles represent maximum voluntary force (MVF) values in Newtons (N) for each participant at baseline (PRE1), after a 20-minute rest period (PRE2), and after occlusion (POST). **B.** Filled circles represent mean H-reflex amplitude as a percentage of maximal M-wave (% M_max_) before (PRE) and after (POST) occlusion for each participant. The mean value at each time point was calculated by taking the average H-reflex amplitude in response to 3 stimulations delivered at least 5 seconds apart during that condition. **C.** Filled circles represent M_max_ amplitude in millivolts (mV) before (PRE) and after (POST) occlusion for each participant. In each figure, the middle horizontal black bar represents the estimated marginal mean obtained from linear mixed modelling with error bars denoting the 95% confidence intervals. *** p < 0.001.

#### H-reflex and M-wave Analysis

As in Experiment 1, after the period of occlusion (mean ± standard deviation of occlusion duration: 21 ± 5 min) soleus H-reflex amplitudes decreased (Χ^2^ = 760.82, p < 0.0001), having been found to be lower at POST (2.70 [2.04, 3.57]% M_max_) compared to PRE (45.40 [34.32, 60.11]% M_max_, p < 0.0001, d = 6.50; Figure 4B). Additionally, there were no significant differences in soleus M_max_ peak-to-peak amplitude across timepoints in Experiment 2 (Χ^2^ = 3.24, p = 0.0718; 4.63 [3.60, 5.79] vs 4.40 [3.40, 5.53] mV, p = 0.0856, d = −0.64; Figure 4C). These results replicate Experiment 1, and indicate that the occlusion protocol likely did not induce changes in peripheral nerve fibre membrane excitability while reducing Ia afferent input to the motor pool.

#### Motor Unit Discharge Characteristics

Peak MU discharge rate differed across time points (Χ^2^_(2)_ = 42.83, p < 0.0001) and between contraction intensities (Χ^2^_(1)_ = 902.45, p < 0.0001), but there was no interaction between time and contraction intensity (Χ^2^_(2)_ = 3.29, p = 0.1928). Peak discharge rate increased at POST (23.5 [21.1, 25.9] pps) compared to PRE1 (22.4 [20.0, 24.8] pps, p < 0.0001, d = 0.36) and PRE2 (21.9 [19.5, 24.3] pps, p < 0.0001, d = 0.52; Figure 5A) across both contraction intensities, however, no difference was found between peak discharge rates at PRE1 and PRE2 (p = 0.0970, d = 0.16).

**Figure 5.**
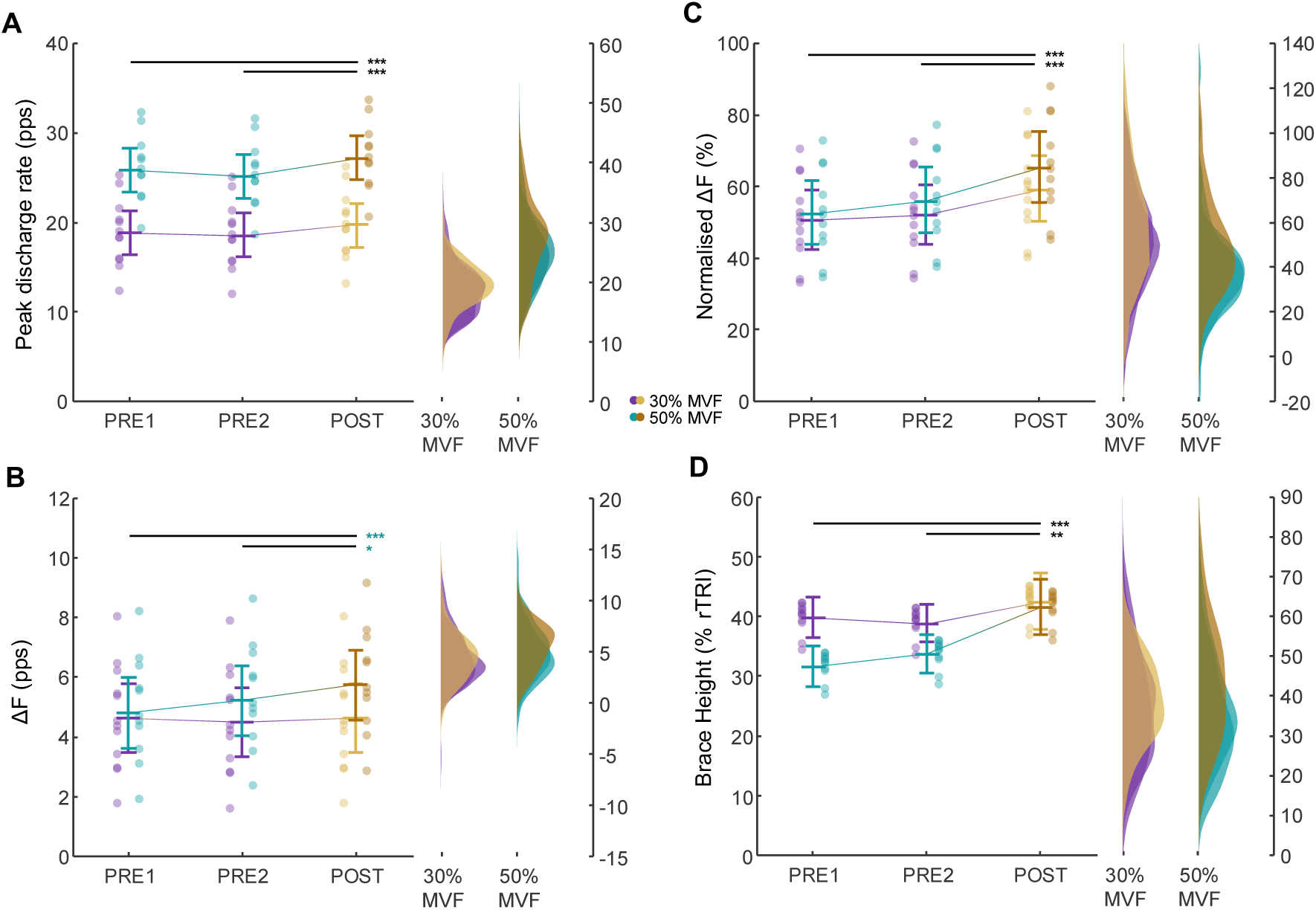
Motor unit discharge characteristics before and after occlusion. **A.** Peak discharge rate in pulses per second (pps), **B.** discharge rate hysteresis (ΔF) in pulses per second (pps), **C.** ΔF normalised to the maximal theoretical hysteresis (%), and **D.** brace height as a percentage of a right triangle (% rTRI) at 30% (purple/gold) and 50% (teal/rust) MVF, at baseline (PRE1), after a 20-minute rest period (PRE2), and after occlusion (POST). Filled circles on the left-hand side of each figure represent individual participant averages over the three conditions at both contraction intensities. The middle horizontal purple/gold and teal/rust bars represent the estimated marginal means obtained from linear mixed modelling with error bars denoting the 95% confidence intervals at 30% and 50% MVF respectively. The kernel density distribution plots on the right-hand side of the figures show the concentration of points from individual motor units at both 30% and 50% MVF across values of peak discharge rate, ΔF, normalised ΔF and brace height respectively. *** p < 0.001, ** p < 0.01, * p < 0.05.

Onset-offset hysteresis changed across experimental time points (Χ^2^_(2)_ = 9.65, p = 0.0080), and differed between contraction intensities (Χ^2^_(1)_ = 19.79, p < 0.0001). There was also a significant interaction between time and contraction intensity (Χ^2^_(2)_ = 10.86, p = 0.0044). During 30% MVF contractions, no difference in onset-offset hysteresis between time points was observed (POST: 4.66 [3.50, 5.82] pps vs PRE1: 4.65 [3.49, 5.81] pps, p = 0.9991, d = 0.00; POST vs PRE2: 4.50 [3.34, 5.66] pps, p = 0.6724, d = 0.10; PRE1 vs PRE2, p = 0.6915, d = 0.10; Figure 5B). However, during 50% MVF contractions, onset-offset hysteresis was greater at POST (5.76 [4.59, 6.94] pps) than at PRE1 (4.82 [3.65, 6.00] pps, p < 0.0001, d = 0.60) and PRE2 (5.24 [4.06, 6.41] pps, p = 0.0376, d = 0.33), but there were no differences in ΔF between PRE1 and PRE2 (p = 0.1088, d = −0.26).

When onset-offset hysteresis was normalised to the maximal theoretical hysteresis for each test-reporter unit pair, values differed across time points (Χ^2^ = 30.80, p < 0.001), but not between contraction intensities (Χ^2^ = 3.62, p = 0.057), and there was no significant interaction between time and intensity (Χ^2^ = 0.91, p = 0.634). For both intensities, normalised ΔF was higher at POST (62.2 [53.5, 71.7]%) compared to PRE1 (51.5 [43.6, 60.1]%, p < 0.0001, d = 0.49; Figure 5C) and PRE2 (54.1 [45.9, 62.8]%, p = 0.0001, d = 0.37), but there was no difference between PRE1 and PRE2 (p = 0.3459, d = −0.12).

Brace height also differed across the three time points (Χ^2^_(2)_ = 15.62, p = 0.0004) and between contraction intensities (Χ^2^_(1)_ = 14.55, p = 0.0001); however, no interaction between time and contraction intensity was observed (Χ^2^_(2)_ = 5.34, p = 0.0692). Across both intensities, brace height was higher at POST (42.0 [38.5, 45.7]%) compared to PRE1 (35.6 [32.9, 38.5]%, p = 0.0004, d = 0.52; Figure 5D) and PRE2 (36.3 [33.7, 39.0]%, p = 0.0014, d = 0.47), but no differences were observed between PRE1 and PRE2 (p = 0.8612, d = −0.05).

## DISCUSSION

Across two experiments, this study examined the effects of reduced afferent input on TA MU discharge characteristics during triangular-shaped isometric dorsiflexion contractions to 30% and 50% MVF. As hypothesised, MU discharge rate and estimates of PICs increased in both experiments. In Experiment 1, there was an increase in peak discharge rate, discharge rate hysteresis and ascending discharge rate non-linearity for ramp contractions performed to 30% of the pre-occlusion MVF. In Experiment 2, when ramp contractions were performed to 30% of each condition’s respective MVF, there was in increase in peak discharge rate and brace height, but not ΔF, post-occlusion. However, once ΔF was normalised to the maximal theoretical hysteresis, an increase was observed from pre- to post-occlusion. During contractions to 50% of each condition’s MVF, there was an increase in peak discharge rate, brace height, ΔF and normalised ΔF, indicating an increase in PIC contribution to the ascending phase modulation and prolongation of MU discharge rate post-occlusion. When taken together, these findings suggest that reduced afferent input via ischaemia may have upregulated the contribution of PICs to MU discharge behaviour during triangular-shaped isometric dorsiflexion contractions, possibly due to reduced reciprocal inhibitory input to the dorsiflexor motor pool.

A prerequisite for interpreting the changes in MU discharge patterns is appreciating the effects of ischaemic occlusion on the sensorimotor system. The almost-complete abolishment of the H-reflex strongly suggests that afferent input from the antagonist muscle was reduced, consistent with evidence suggesting preferential vulnerability of conduction in myelinated sensory axons to ischaemia (Hofmeijer et al., 2013; Parry & Brown, 1982). Conversely, M_max_ remained relatively unchanged after occlusion compared to before, confirming that motor axon conduction was largely retained, consistent with prior work (Hofmeijer et al., 2013; Lorentzen et al., 2018; Rasul et al., 2022). Nevertheless, the exclusion of three participants from the analysis of Experiment 1 due to the inability to reach the target forces post-occlusion, suggested motor function was impaired by occlusion, a finding that was later confirmed in Experiment 2. Although small efferent drive impairment cannot be excluded as a possible cause, it is likely that the reduced contractile function after occlusion is the result of a partial block of Ia afferent-mediated agonist proprioceptive feedback, leading to reduced excitatory synaptic input to the agonist motor pool. As such, and with an appreciation of the crude nature of the ischaemic occlusion employed in this study, we primarily interpret the observed changes in MU discharge patterns as a function of reduced afferent input, likely, at least in part, mediated by reduced disynaptic reciprocal inhibitory drive to the agonist motoneurons as evidenced by reduced H-reflex (Crone et al., 1987).

### Effects of ischaemia on motor unit discharge during voluntary contractions to the same absolute force

In Experiment 1, an increase in peak discharge rate was observed post-occlusion compared to pre-, and this was accompanied by greater MU discharge rate hysteresis and less linear ascending MU discharge patterns. Importantly in this experiment, these changes post-occlusion were observed during contractions performed to the same *absolute* force. As indicated in the previous section, some reduction in contractile function likely occurred as a result of occlusion. Therefore, the greater peak MU discharge rate and hysteresis (ΔF) is difficult to attribute solely to the possible reduction in reciprocal inhibition and the resultant upregulation of PICs (Heckman et al., 2005) as both of these metrics are known to increase with greater contraction force (Goodlich et al., 2023; Mackay et al., 2023; Orssatto et al., 2021; Škarabot et al., 2025). Indeed, when normalised to the maximal theoretical hysteresis of a test-reporter unit pair, ΔF was found to be unchanged following occlusion, suggesting that the additional synaptic input to achieve higher relative forces post-occlusion could have masked its effects.

Nevertheless, the increase in the ascending MU discharge non-linearity suggests that, at least in part, the changes in peak MU discharge rate and hysteresis could be attributable to upregulation of PICs, likely due to reduction in reciprocal inhibition via the afferent block. That is, if the observed effects were solely a function of greater absolute force levels, one would expect the ascending MU discharge rates to linearise as shown previously (Škarabot et al., 2025). This result therefore suggests that the afferent block via ischaemia led to a greater relative contribution of PICs to the ascending discharge rate (Beauchamp et al., 2023; Chardon et al., 2024).

### Effects of ischaemia on motor unit discharge during voluntary contractions to the same relative force

Owing to the observation of the possible loss of contractile function in Experiment 1, Experiment 2 involved contractions performed to the same *relative* force level. That is, maximal voluntary force was reassessed at each time point to account for decrements in contractile function, which was evident following occlusion, likely due to Ia-mediated reduction in excitatory synaptic input to the agonist motor pool. Additionally, because of different requirements for neuromodulation and modulation of excitation-inhibition coupling (Škarabot et al., 2025), and the recruitment of additional, higher-threshold MUs, we wanted to assess whether potential effects of the loss of afferent input on MU discharge characteristics are contraction intensity-dependent.

Similar to Experiment 1, we showed greater MU peak discharge rate following occlusion and this did not depend on contraction intensity. We also observed greater ascending MU discharge rate non-linearity, regardless of contraction intensity, consistent with greater relative contribution of PICs to the ascending MU discharge rate (Beauchamp et al., 2023; Chardon et al., 2024). Onset-offset hysteresis, on the other hand, was observed to increase post-occlusion during contractions to 50% MVF, but not 30% MVF, indicating there may be a contraction intensity-dependent response. However, when ΔF was normalised to the maximal theoretical hysteresis of the test-reporter unit pair, an increase was found post-occlusion at both 30% and 50% MVF. The discrepancy in the absolute and relative ΔF changes likely stems from floor effects of the ΔF calculation where long descending discharge durations might have been present at baseline in the 30% MVF condition, making detection of additional changes in hysteresis less likely. Taken together, these results suggest that a reduction in afferent input via occlusion upregulates PIC contribution to MU discharge rate, possibly due to decreased reciprocal inhibition. As such these results provide additional evidence for the possible role of reciprocal inhibition in sculpting the contributions of PICs to MU discharge patterns in humans (Gomes et al., 2024; Mesquita et al., 2022; Orssatto et al., 2022; Pearcey et al., 2022), but represent the first experimental effects of *reduced*, rather than *increased* reciprocal inhibition on PICs, which appear to be largely uniform across contraction intensities.

### Limitations and alternative explanations

Although the results strongly point to the upregulation of PIC contribution to MU discharge rate via decreased reciprocal inhibition as the mechanism for observed changes in MU discharge characteristics in our experiments, reduction in afferent input via occlusion is a relatively crude technique, and thus several potential contributing factors exist. First, we cannot exclude the possibility that the observed loss of afferent input resulted in an increase in descending drive to the agonist motor pool, in addition to the changes in reciprocal inhibitory input to the agonist motoneuron. However, this possibility is less likely due to the increase in the ascending discharge rate non-linearity, which typically decreases with greater excitation (Škarabot et al., 2025). Future studies could employ transcranial magnetic stimulation to assess corticospinal excitability following ischaemic block of afferent input to assess the relative contribution of changes in descending drive to the observed MU discharge characteristics. Second, the occlusion could have stimulated group III and IV afferents, which have been shown to be sensitive to ischaemia (Jankowski et al., 2013; Light et al., 2008; Pollak et al., 2014) and are known to modulate H-reflex responses (Taylor et al., 2016). The contribution of group III and IV afferents is further complicated by the performance of contraction under occlusion, conditions under which these afferents are likely to be excited due to metabolite build up (Li et al., 2003; Rybicki et al., 1985), which may have affected MU discharge characteristics (Taylor et al., 2016). We intended to minimise these effects by performing only a single or, in the case of Experiment 2, two contractions. Still, further studies could include multiple contractions (Lowe et al., 2023) to assess the extent to which MU discharge patterns characterised in the present study would be modulated by the likely additional excitation of group III and IV afferents. Finally, we cannot fully exclude potentially subtle modifications in motor axon conduction that remained undetectable in the present study (i.e., no changes in maximal M-wave following occlusion). However, Experiment 2 was designed to control for the most obvious effect of this potential modification, and the persistence of changes in MU discharge characteristics at matched forces, consistent with our hypothesis, suggests any potential intrinsic changes to motor axon conduction were minimal.

## Conclusion

In conclusion, we showed that estimates of PICs appear to increase with reduced afferent feedback resulting from ischaemic nerve block. Specifically, we showed that peak discharge rate, onset-offset hysteresis and non-linearity in ascending discharge rate increased post-occlusion during isometric dorsiflexion contractions performed to the same *absolute* contraction intensity. When force was expressed relative to the reduced post-occlusion MVF, peak discharge rate, discharge rate hysteresis, and non-linearity in ascending discharge rate still increased post-occlusion, and these observations were apparent at both 30% and 50% MVF. These findings suggest that reduced afferent input, likely resulting in a reduction in reciprocal inhibitory input, leads to an upregulation of estimates of PICs. This has implications for cases of neuromuscular disease resulting in deafferentation or loss of afferent input.

## Notes

### Competing Interest Statement

The authors have declared no competing interest.

### Funding

- Ministry for Education, Sport, Youth, Research and Innovation, MT

